# The mediodorsal thalamus supports adaptive responding based on stimulus-outcome associations

**DOI:** 10.1101/2022.09.12.507585

**Authors:** Sarah Morceau, Angélique Faugère, Etienne Coutureau, Mathieu Wolff

## Abstract

The ability to engage into flexible behaviors is crucial in dynamic environments. We recently showed that in addition to the well described role of the orbitofrontal cortex (OFC), its thalamic input from the submedius thalamic nucleus (Sub) also contributes to adaptive responding during Pavlovian degradation. In the present study, we examined the role of the mediodorsal thalamus (MD) which is the other main thalamic input to the OFC. To this end, we assessed the effect of both pre- and post-training MD lesions in rats performing a Pavlovian contingency degradation task. Pre-training lesions mildly impeded the establishment of stimulus-outcome associations during the initial training of Pavlovian conditioning without interfering with Pavlovian degradation training when the sensory feedback provided by the outcome rewards were available to animals. However, we found that both pre- and post-training MD lesions produced a selective impairment during a test conducted under extinction conditions, during which only current mental representation could guide behavior. Altogether, these data suggest a role for the MD in the successful encoding and representation of Pavlovian associations.

## Introduction

Representing and maintaining accurate knowledge about the environment is critical for the survival of any organism. In natural environments, the relevance of incoming signals may vary over time, which prompts the need to regularly track the current predictive value of environmental cues. Pavlovian conditioning paradigms provide a rich framework to examine how predictive cues are used by animals to adjust behavioural output. In particular, degrading the contingency between a stimulus and its associated outcome provides an ideal avenue to examine how animals can adaptively adjust behavior in response to a cue that was previously relevant but that is now no longer reliable (Delamater, 1995). This process, called Pavlovian degradation has proven to be quite effective to highlight the neural bases of adaptive behaviors. Importantly, highly homologous neuronal circuits appear to support these abilities in mammals (Balleine and O’Doherty, 2010), with an important role for prefrontal regions to control behaviour and adapt it to current circumstances.

The orbitofrontal cortex (OFC) in particular appears as a core area for the encoding and updating of Pavlovian stimulus-outcome associations (Ostlund and Balleine, 2007). For instance, OFC lesions typically abolish the capacity to update these associations (Ostlund and Balleine, 2007); Alcaraz et al., 2015) and neuronal activity within the OFC encode both the value and the identity of predicted outcomes (Howard and Kahnt, 2021, 2018; Stalnaker et al., 2018). As a highly integrative hub, the OFC connects to multiple brain regions especially at the subcortical level. Thalamic nuclei have recently emerged as essential partners for the cortical stage to support cognitive functions (Wolff et al., 2015; Wolff & Vann, 2019; Pergola et al., 2018; Rikhye et al., 2018; Perry et al., 2021; Wolff et al., 2021). Interestingly, the OFC is the target from convergent inputs arising from two distinct thalamic nuclei, the submedius thalamic nucleus (Sub) and the mediodorsal thalamus (MD) (Alcaraz et al., 2015; Alcaraz et al., 2016; Murphy & Deutch, 2018; Kuramoto et al., 2017a, 2017b). Previously, we identified the submedius thalamus as critical for updating stimulus-outcome associations (Alcaraz et al., 2015) and we also confirmed that functional interactions between the OFC and the Sub are necessary for adaptive responding (Fresno et al., 2019). While few evidence suggest that the MD may also guide behaviour based on predictive cues (Pickens, 2008; Ostlund & Balleine, 2008), the role of this region has not been examined within the same experimental framework, known to be reliant on OFC functions (Alcaraz et al., 2015).

The present study aimed at directly assessing the role of the MD in the flexible use of predictive cues, in a Pavlovian degradation protocol previously shown to be sensitive to both OFC and Sub damage. To probe MD functions, we initially performed pre-training MD neurotoxic lesions in rats subsequently submitted to Pavlovian degradation. During degradation training per se, rats harbouring MD lesions were able to adapt to the new Pavlovian contingencies, as did Sham rats, despite a modest impairment in initial Pavlovian conditioning. But during a test conducted under extinction condition they were unable to maintain differential responding for the two predictive cues. To rule out a possible confound with initial acquisition which was affected by the lesion, we then adopted a post-training lesion approach and we fully replicated these findings: rats with MD lesions behaved normally during degradation training but exhibited a selective impairment when the sensory feedback provided by the food outcome was omitted during a test conducted under extinction conditions. Altogether, these data support the idea that the MD is important to guide behavior based on current mental representations (Wolff and Vann, 2019).

## Methods

### Animals and housing conditions

Forty-four male Long-Evans rats weighting 250g to 300g at surgery were obtained from the Centre d’Elevage Janvier (France). Rats were initially housed in pairs and accustomed to the laboratory facility for two weeks before the beginning of the experiments. Environmental enrichment was provided by a polycarbonate tubing element in accordance with current French (Council directive 2013–118, February 1, 2013) and European (directive 2010–63, September 22, 2010, European Community) laws and policies regarding animal experiments. The temperature was maintained at 21 ± 1°C with lights on from 7 a.m. to 7 p.m. For all behavioral experiments rats were food restricted to be around 90-95% of their initial body weight.

### Surgery

Rats were anesthetized with 4% isoflurane and placed in a stereotaxic frame with atraumatic ear bars, in a flat skull position. During the surgery, isoflurane was maintained at 1.5 – 2%. Bilateral neurotoxic lesions were made using 20μg/μl NMDA microinjections (Sigma-Aldrich). Glass micropipettes (outside diameter ∼ 50μm) connected by polyethylene tubing to a Picospritzer (General Valve) were used for pressure injections. One lesion per side as MD lesions were made with one lesion per side as follows: AP, -2.7; laterality, **±** 0.8; DV, -5.0mm from dura. Each site was injected with 0.15μl of NMDA. The Sham groups received similar surgery except that the micropipette was inserted only in the cortex with no injection as follows: AP: -2.7; laterality, **±** 0.7; DV from dura -2.0mm. For all groups, the micropipette was left in place 3min after injection before a slow retraction. Rats were given 8-10 days of recovery before behavioral testing.

### Behavioral experiments

#### Behavioral apparatus

Eight identical conditioning chambers (40cm wide X 30cm deep X 35cm high; Imetronic) were used for behavioral experiments. Each chamber was located inside a sound and light attenuating wooden chamber (74 × 46 × 50 cm). Each of them had a ventilation fan that produce a background noise of 55dB and 4 LEDs on the ceiling for illumination. Chambers had two opaque panels on each side (right and left) and a stainless-steel grid floor (rod diameter, 0.5 cm; inter-rod distance, 1.5 cm). A magazine (6 × 4.5 × 4.5 cm), placed in the middle of the left wall, could collect either food pellets (45 mg; F0165, Bio-Serv) or sucrose pellets (45 mg; 1811251, Bio-Serv) from dispensers located outside the operant chamber. Speakers in each chamber provided either a 3 kHz Tone or a 10 Hz Clicker auditory stimulus, both produced by the activation of a mechanical relay. The magazine was equipped with infrared cells to detect the animal’s visits. A personal computer connected to the conditioning chambers enabled to control the equipment and record the data (Poly Software, Imetronic).

#### Pavlovian contingency degradation

To examine the functional contribution of the MD in flexible outcome-guided behaviors, we aimed to focus on the ability to encode and update Pavlovian contingencies. To do so, an initial appetitive Pavlovian conditioning is conducted, during which rats were required to learn two distinct stimulus-outcome associations. Once the Pavlovian associations were reliably established, the contingency between one of the conditional stimuli (CS) and its outcome was selectively degraded so that the CS no longer reliably predicted the reward. Overall, there are three distinct phases of behavioral testing: Pavlovian conditioning, contingency degradation training and finally a test conducted under extinction conditions.

### Pavlovian conditioning

The conditioning phase consisted in eight forty minutes daily sessions during which rats learned that each predictive auditory cue was associated with the delivery of a particular outcome (i.e. grain or sugar pellets). For each session, each of the two CS (either the Tone or the Clicker) was presented 15 times consecutively. Each CS was presented for twenty seconds during which two samples of the associated reward were delivered. CS were separated by an average intertrial interval (ITI) of 60s. Specific associations were counterbalanced within groups (half have grain pellets associated with the tone and sucrose pellets with the clicker and half the alternate associations) for a total of 30 grain and 30 sucrose pellets delivered per daily session.

### Pavlovian contingency degradation

Following Pavlovian conditioning, all rats were given six additional daily sessions. The only difference with Pavlovian conditioning was that one stimulus-outcome association was degraded, as animals now had an equal probability to get the food reward during CS presentation or during the ITI; the overall number of rewards was thus maintained. The nondegraded CS and its associated outcome were presented with the same contingencies as that used for the Pavlovian conditioning phase. As a result, rats should learn that this particular CS (degraded CS) has become less reliable to predict the delivery of the outcome and they are expected to diminish responding to that CS. All stimulus-outcome associations and associated contingency schedules (degraded *versus* nondegraded) were counterbalanced across rats and lesion groups.

### Test without rewards (in extinction)

One day after the last session of contingency degradation, rats underwent a final test under extinction conditions. This test consisted of four presentations of each CS presented in alternation (duration of the CS: 20s, duration of the ITI: 60s, total duration of the test: 11.40min). No reward was delivered during this test, preventing thus the animal to benefit from the sensory feedback of the rewards. Therefore, on this occasion, the animal’s responding is only guided by its current mental representations.

### Histology

Animals received a lethal dose of sodium pentobarbital and were perfused transcardially with 150 mL of saline followed by 150mL of 10% formaldehyde. The sections of the MD were cut with a vibratome at 60μm. Sections were then collected onto gelatin-coated slides and dried before being stained with thionine. Histological analysis of the lesions was performed under the microscope by two experimenters (MW and SM) blind to lesion conditions.

### Data analysis

The data were submitted to ANOVAs on StatView software (SAS Institute) with Lesion (Sham/MD), as between subject factor, Period (CS, ITI), Degradation (Degraded/NonDegraded), and Session as repeated measures. The dependent measure of interest was the average frequency of magazine visits during the CS (average number of visits per minute for acquisition and % of baseline performance for degradation training and the final test; baseline performance corresponding to the final session of Pavlovian conditioning), or preceding the CS (ITI) for initial conditioning. For the test conducting under extinction conditions, we focused our analysis on the first whole 20sec CS presentation for both degraded and non-degraded conditions to minimize any confound with extinction. To demonstrate that responding was stable during the presentation of these first stimuli we computed responding across four 5sec bins. To control for any potential nonspecific effect of the lesion we also computed a score by subtracting to the number of visits recorded during stimuli presentation, that recorded during the immediately preceding 20sec Pre-CS period. The α value for rejection of the null hypothesis was 0.05 throughout.

## Results

### Histology

Overall, MD lesions (Figure 1A) were highly similar to previous works (Alcaraz et al., 2016; Wolff et al., 2015), showing substantial damage to the whole region, including the central, medial and lateral segments along the full extent of the anteroposterior axis (Figure 1B). Lesions tended to encroach onto adjacent thalamic nuclei and especially onto the intralaminar nuclei, more particularly the centromedian and paracentral thalamic nuclei, while the centrolateral nuclei was only marginally affected. Additional damage to the paraventricular thalamic nucleus was also apparent in several cases. This additional damage to surrounding thalamic nuclei was not associated with any specific behavioural profile and the magnitude of the deficit was similar across all included individuals. For experiment 1, one MD rat was discarded from the behavioral analyses as it exhibited only minimal damage. In addition, two rats died after surgery (1 Sham and 1 MD). For experiment 2, one lesioned rat was discarded because the lesion was too posterior. Thus, the final groups for experiments 1 and 2 were therefore as follows: Sham Pre: n = 9; MD Pre: n = 10; Sham Post: n =10; MD Post: n = 11.

**Figure 1.**
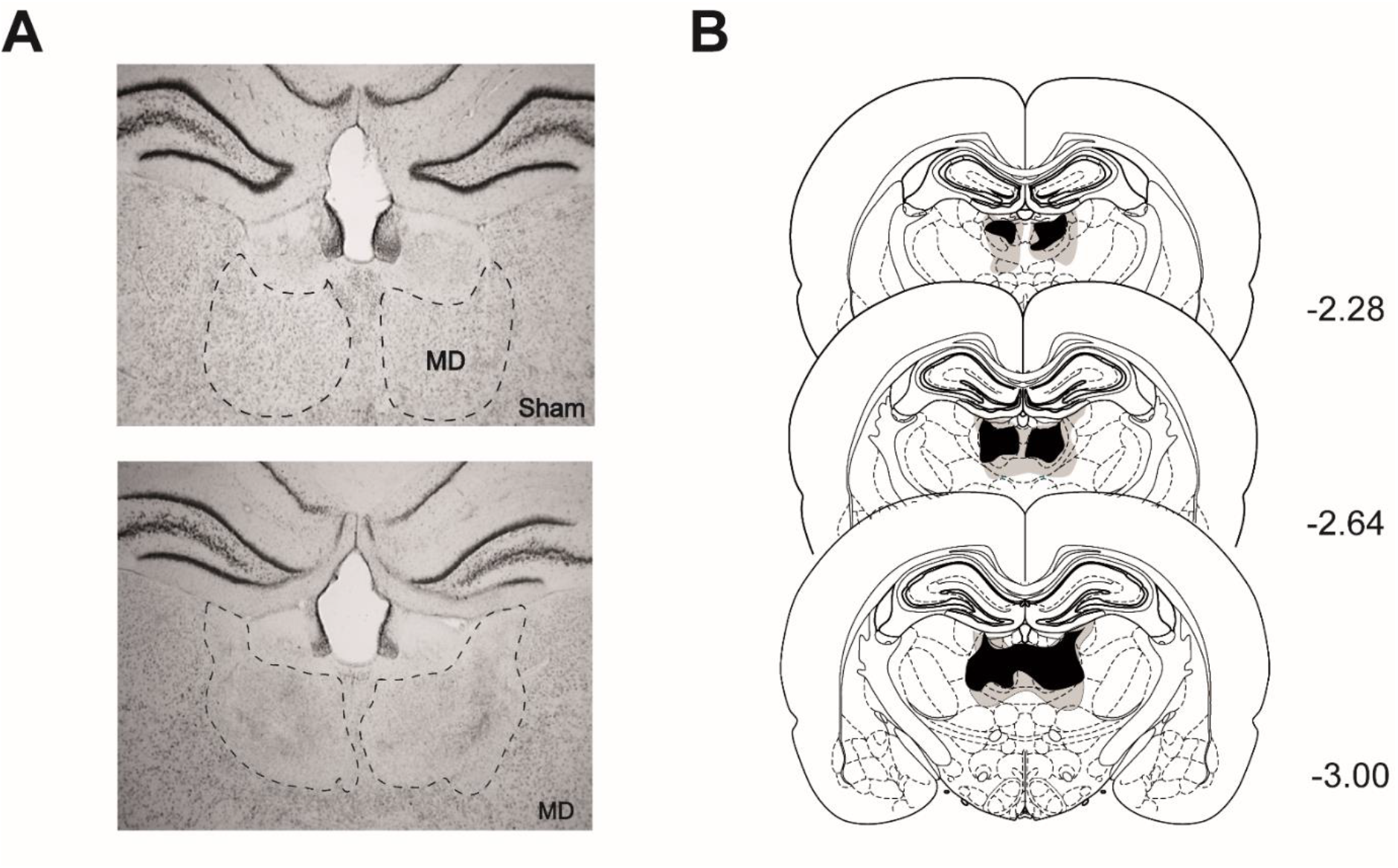
Histology. **A**. Representative photomicrographs of the MD (dashed line) in Sham (top) and lesioned rats (bottom). **B**. Schematic representation of the included largest (gray) and smallest (black) MD lesion at three different levels of the anteroposterior axis (indicted in millimetres relative to bregma), Thionine protein stain.

### Experiment 1: Pavlovian contingency degradation in pre-training lesioned rats

#### Pavlovian training

Figure 2 shows the average rate of visits to the magazine during both CS presentation and the period preceding the stimulus (ITI). All rats preferentially visited the magazine during the CS, indicating that they progressively learned Pavlovian associations. These observations were supported by highly significant effects of Period (F_(1,17)_ = 89.00 P < 0.0001), Session (F_(7,119)_ = 28.51, P < 0.0001) and of the Session X Period interaction (F_(7,119)_ = 11.69, P < 0.0001). Overall, responding appeared to be somewhat lower in the MD group. The analyses indeed confirmed significant effects of Lesion (F_(1,17)_ = 5.62, P = 0.0299) and of the Lesion X Session (F_(7,119)_ = 6.62, P < 0.0001), Lesion X Period (F_(1,17)_ = 4.76, P = 0.0435), and Lesion X Period X Session (F_(7,119)_ = 2.78, P = 0. 0104) interactions. In view of these differences, we conducted supplemental analyses in each group separately, which confirmed that Period reached significance for both MD (F_(1,9)_ = 43.09, P = 0.0001) and Sham (F_(1,8)_ = 45.77, P = 0.0001) rats, and so did the Period X Session interaction (MD, F_(7,63)_ = 2.41, P = 0.0301; Sham, F_(7,56)_ = 10.28, P < 0.0001). Pre-training lesion of MD did thus not prevent Pavlovian conditioning but reduced responding was evident in MD rats.

**Figure 2.**
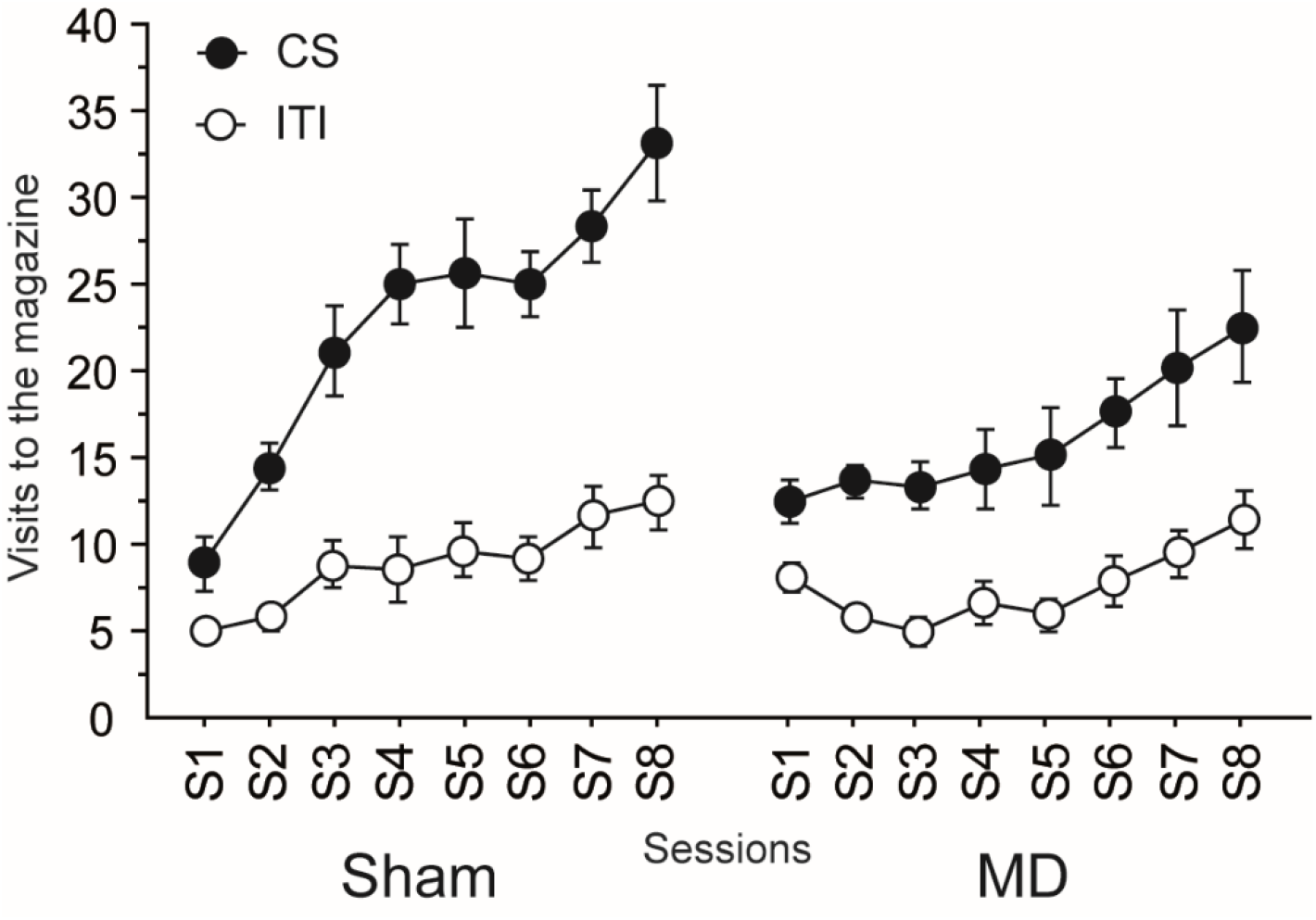
Pre-training MD lesions: acquisition of Pavlovian associations. Pavlovian conditioning, number of visits to the magazine (per minute) during the CS and the ITI.

#### Pavlovian contingency degradation

Figure 3A shows the rate of visits to the magazine during Pavlovian contingency degradation for the cues with either a degraded or nondegraded predictive value. At this occasion, all rats displayed adaptive responding, as responding to the stimulus corresponding to the degraded Pavlovian association progressively decreased. These observations were supported by the main effect of Degradation (F_(1,17)_ = 20.87, P = 0.0003) and a significant Degradation X Session interaction (F_(5,85)_ = 4.89, P = 0.0006). Highly similar responding across groups was evident throughout this degradation phase as neither the factor Lesion (F_(1,17)_ = 0.71, P = 0.4110) nor the Lesion X Degradation (F_(1,17)_ = 0.04, P = 0.8496) or the Lesion X Degradation X Session (F_(5,85)_= 0.20, P = 0.9621) interaction reached significance. All rats thus appeared to adapt their behavior to the new Pavlovian contingencies when food outcomes are available to the animals despite the mild effect of MD lesion during the initial acquisition.

**Figure 3.**
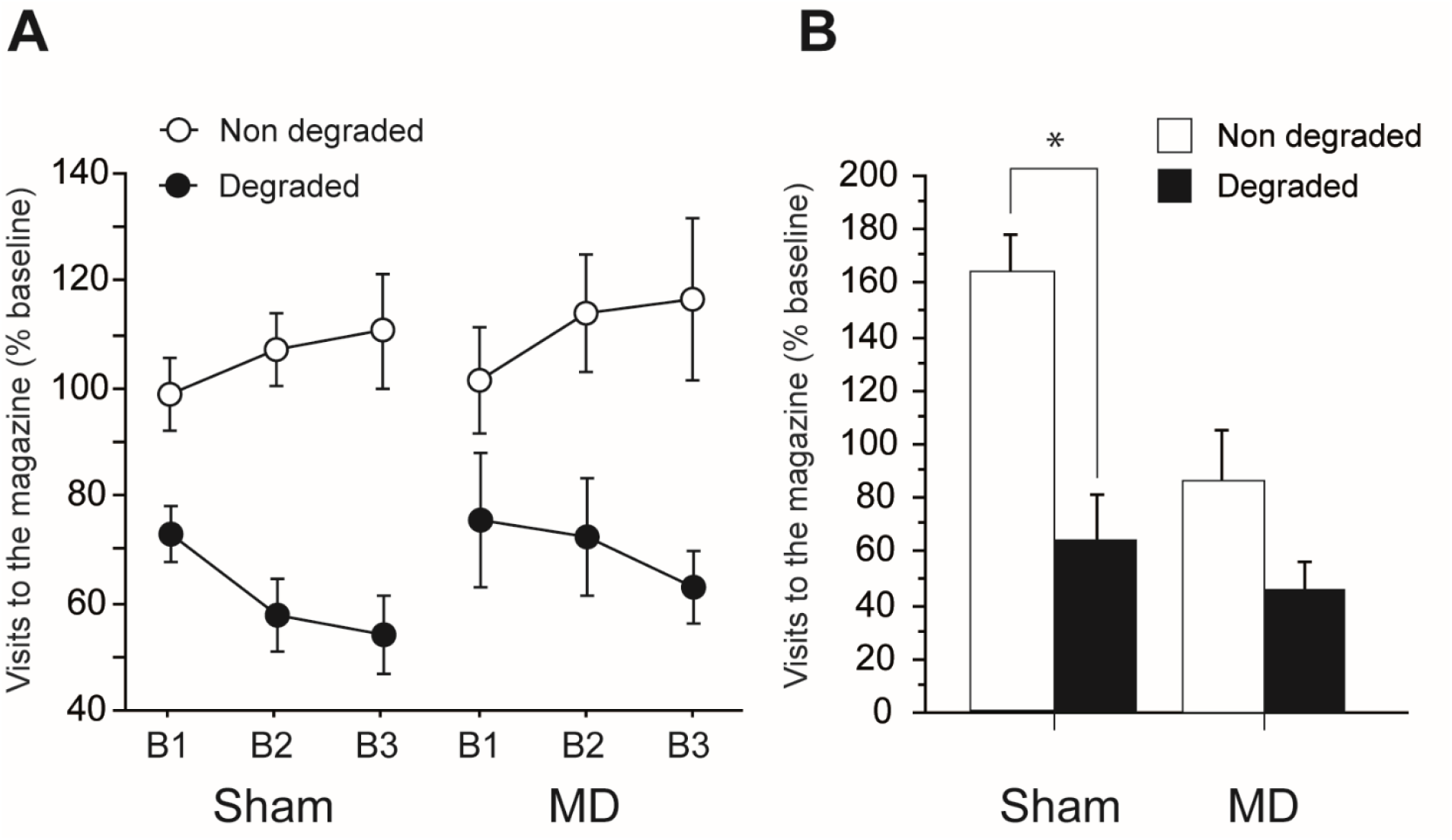
Pre-training MD lesions: Pavlovian contingency degradation. Magazine visits expressed relative to the last session of acquisition (% baseline) during **(A)** degradation training and **(B)** the test conducted under extinction conditions. Results are shown for the nondegraded (white) as well as the degraded (black) contingencies for Sham (left) and MD (right) groups. Data are expressed as mean ± SEM ; * P < 0.05.

#### Test conducted under extinction conditions

During presentation of the first stimuli at test, rats from both groups expressed stable responding across the whole 20s duration (5sec Bloc, 5sec Bloc X Lesion, Fs < 1).

Figure 3B displays the rate of visits to the magazine during the test conducted under extinction conditions. The Sham group continued to exhibit differential responding by visiting more frequently the magazine during the presentation of the CS with nondegraded predictive value. By comparison, MD rats now exhibited reduced responding for both CS. The critical Degradation X Lesion interaction indeed reached significance (F_(1,17)_ = 4.84, P = 0.0419), consistent with the notion that only Sham rats maintained differential responding during the test. Further analyses indeed confirmed that a main effect of Degradation was highly significant for Sham rats (F_(1,8)_ = 31.64, P = 0.0005) but that it only approached significance for MD rats (F_(1,9)_ = 4.29, P = 0.0683).

To further confirm the selectivity of these results, we also computed a difference score by subtracting the number of visits during a 20sec PreCS period to that during the immediately following CS, thus controlling for baseline activity in-between CS presentations. This analysis produced highly significant effects of Lesion (F_(1,17)_ = 19.67, P = 0.0004) and Degradation (F_(1,17)_= 12.46, P = 0.0026) and also a highly significant Lesion X Degradation interaction (F_(1,17)_ = 7.34, P = 0.0149). For this parameter, the effect of Degradation was found to be significant for Sham (F_(1,8)_ = 11.34, P = 0.0098, but not MD rats (F<1), thus fully confirming the initial analysis.

Altogether, these data suggest that MD rats may be generalizing the adaptive response during the test conducted under extinction condition, during which the sensory feedback provided by the food outcome cannot guide behavior. Since the impairment evident during the initial conditioning phase may account to some account for this result, it provided the incentive to examine the impact of post-training MD lesion in the next experiment.

### Experiment 2: Pavlovian contingency degradation in post-training lesioned rats

#### Pavlovian training

During initial Pavlovian conditioning, all rats exhibited higher rate of visits to the magazine during the CS compared to the ITI across sessions (Figure 4). The analyses indeed showed highly significant effects of Period (F_(1,19)_ = 126.78, P < 0.0001), Session (F_(7,133)_ = 49.83, P < 0.0001) and of the Period X Session interaction (F_(7,133)_ = 7.84, P < 0.0001) exactly as before. Before proceeding to post-training lesions, we built two equivalent groups that were matched for presurgery performance as confirmed by the analyses (Lesion (F_(1,19)_ = 1.96, P = 0.1778; Lesion X Period, Lesion X Session, Lesion X Period X Session, Fs <1). Thus, prior to surgery all rats learned Pavlovian associations in a similar way.

**Figure 4.**
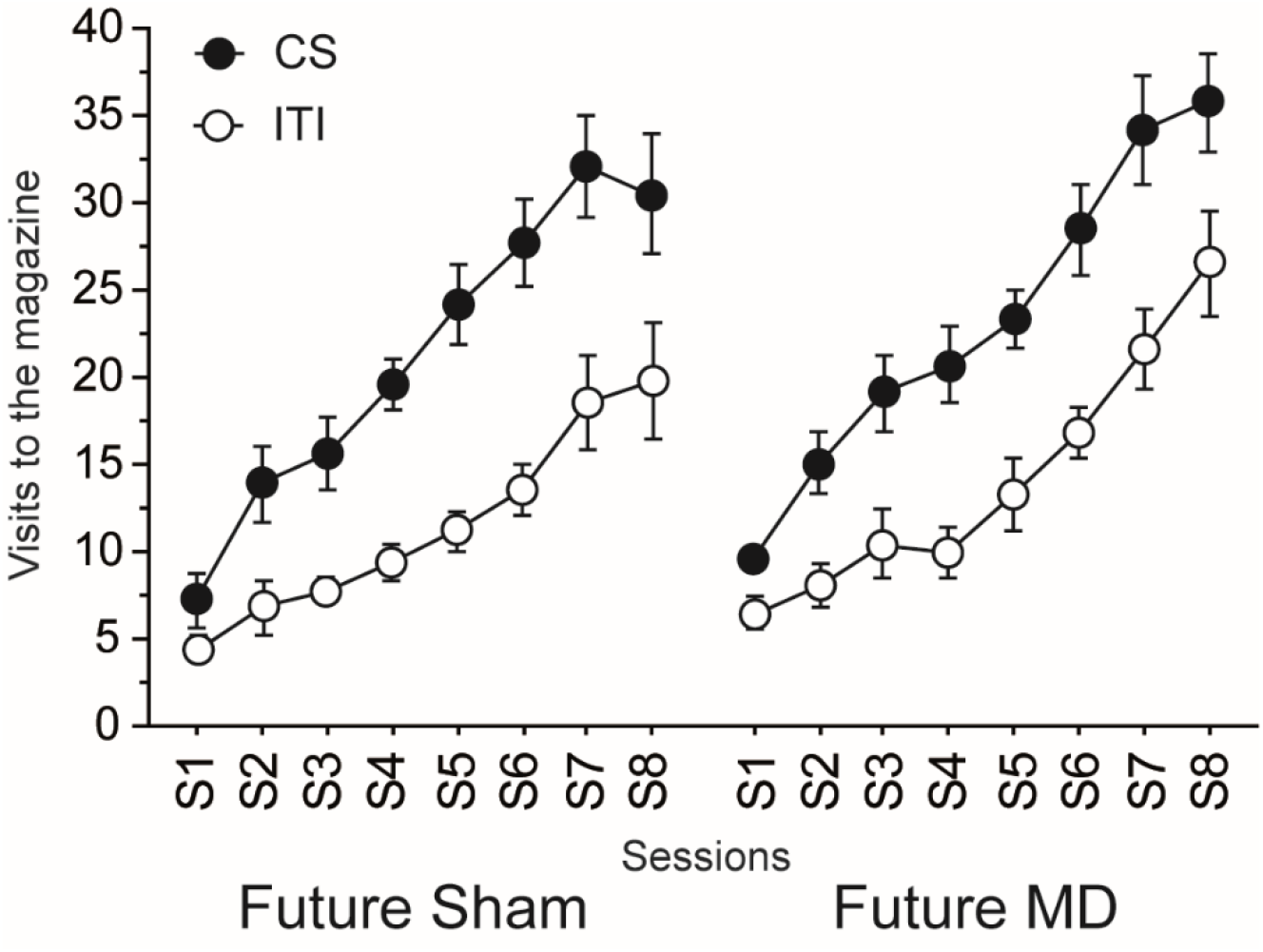
Post-training MD lesions: acquisition of Pavlovian associations in the future Sham and MD groups (surgery was performed after this stage). Pavlovian conditioning, number of visits to the magazine (per minute) during the CS and the ITI.

#### Pavlovian contingency degradation

During Pavlovian contingency degradation, all rats displayed adaptive responding, by progressively reducing their response for the stimulus for which the Pavlovian association is altered (Figure 5A). The main effect of Degradation (F_(1,19)_ = 5.19, P = 0.0345), Session (F_(5,95)_ = 2.85, P = 0.0191) and the Degradation X Session interaction (F_(5,95)_ = 2.75, P = 0.0231) reached significance, thus confirming this observation. As before, we could not detect any difference between groups throughout this degradation phase as neither Lesion (F_(1,19)_ = 1.56, P = 0.23) nor the Lesion X Degradation (F<1) or Lesion X Degradation X Session (F_(5,95)_ = 1.16, P = 0.3351) interactions reached significance. As before for pre-training lesions (experiment 1), all rats adapted their response to the new Pavlovian associations when food outcomes were available.

**Figure 5.**
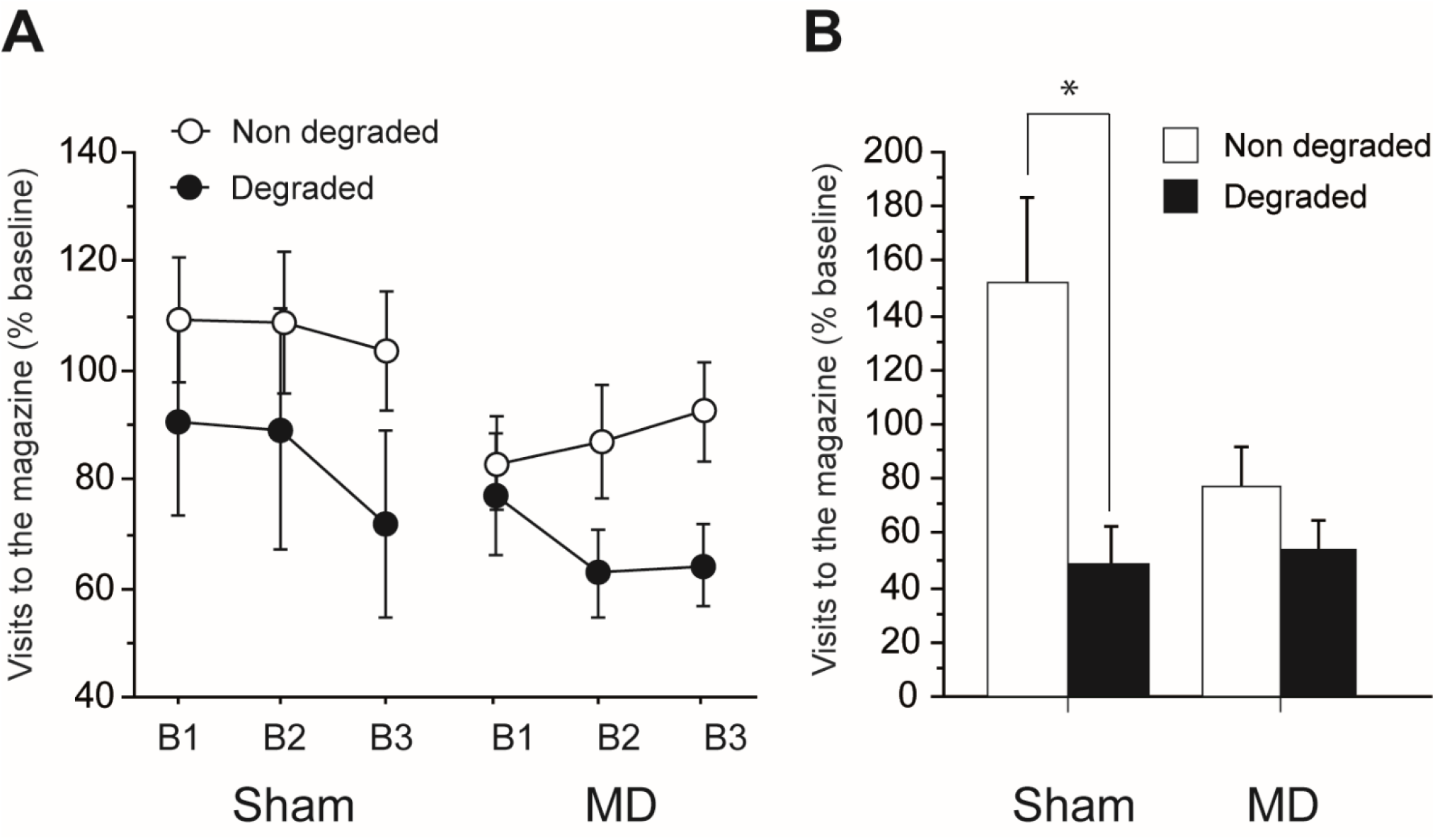
Post-training MD lesions: Pavlovian contingency degradation. Magazine visits expressed relative to the last session of acquisition (% baseline) during **(A)** degradation training and **(B)** the test conducted under extinction conditions. Results are shown for the nondegraded (white) as well as the degraded (black) contingencies for Sham (left) and MD (right) groups. Data are expressed as mean ± SEM ; * P < 0.05.

#### Test conducted under extinction conditions

During the test, the Sham group maintained differential responding for the degraded and the non-degraded stimulus (Figure 5B). Unlike Sham rats, MD rats again exhibited reduced responding for both stimuli. The main effect of Degradation was significant (F_(1,19)_ = 14.08, P = 0.0014) but the main effect of Lesion was not (F_(1,19)_ = 2.69, P = 0.1176). Importantly, the critical Degradation X Lesion reached significance (F_(1,19)_ = 5.74, P = 0.0270) confirming differential profiles in Sham *versus* MD rats. More specifically, the main effect of Degradation reached significance for the Sham (F_(1,9)_ = 11.66, P = 0.0077) but not the MD (F_(1,10)_ = 1.90, P = 0.1982).

A similar conclusion was reached when analyzing the difference score. While the analysis revealed that the effect of Lesion only approached significance (F_(1,19)_= 3.52, P = 0.0761), the main effect of Degradation reached significance (F_(1,19)_= 14.81, P = 0.0011) and the critical Lesion X Degradation interaction was also significant (F_(1,19)_= 5.15, P = 0.0351). As in the first experiment, the effect of Degradation was significant for the Sham (F_(1,9)_ = 19.37, P = 0.0017) but not the MD group (F_(1,10)_ = 1.22, P = 0.2945) highlighting again the consistent results between the first and the second experiments.

This pattern of results thus mirrors that observed for experiment 1, with animals that were intact during the initial Pavlovian conditioning phase. Overall, degradation training proper was not affected by MD lesions even when the initial acquisition is impacted by MD lesion (experiment 1) but both pre- and post-training MD lesions produced a severe impairment during the test under extinction conditions, indicating that rats harboring MD lesions are unable to guide their behavior based on current stimulus-outcome associations if no sensory feedback is available.

## Discussion

Thalamic nuclei provide important inputs to multiple subdivisions of the prefrontal cortex which may account for the important integrative functions of this cortical region. Thalamocortical projections indeed contact not only the OFC but also the medial prefrontal cortex (mPFC) (Alcaraz et al., 2016; Groenewegen, 1988; Phillips et al., 2019) In the present work, we examined the possible role of the MD in flexible responding when stimulus-outcome associations guide behavior, a function previously demonstrated to be reliant on OFC functions. We found that both pre- and post-training MD lesions resulted in an inability to maintain flexible responding when the sensory feedback provided by the food outcome was not available to the animals. This convergent pattern of results from two independent experiments is consistent with the view that the MD plays a critical role in guiding behaviour by relying on current mental representations.

The present set of results indicate that during degradation training *per se*, rats with MD lesions appeared to develop adaptive differential responding for the two cues, in a manner that was largely equivalent to that of the Sham group and it was the case for both experiments. This clearly indicates that MD lesions did not prevent rats from identifying a change in cue-outcome contingencies or from engaging into successful adaptive strategies. While the former point is consistent with previous findings (Ostlund and Balleine, 2008), the latter differs from that earlier study during which Pavlovian degradation training was found to be impaired by MD lesions even when outcomes were delivered. This discrepancy may result from methodological differences: while rats in the Ostlund & Balleine study went through multiple phases of testing involving learning both instrumental and Pavlovian associations, the present study was conducted with a pure Pavlovian framework. As a result, it is possible that rats were facing a more serious behavioral challenge in the initial study, which may be more sensitive to MD dysfunction. The ability of rats with MD lesion to cope with the Pavlovian degradation procedure when outcomes were available also differs from previous works on instrumental degradation (Corbit et al., 2003; Parnaudeau et al., 2015 but see Alcaraz et al., 2018) and suggests a possibly more limited role of the MD in cue-guided choices, except when instrumental and Pavlovian contingencies are mixed (Alcaraz et al., 2016; Ostlund and Balleine, 2008), possibly suggesting a more prominent role of this region in complex behavioural settings (Mukherjee et al., 2021). However, when the sensory feedback provided by food outcomes was removed during the tests conducted under extinction conditions, a large and specific deficit was consistently apparent in rats with MD damage for both experiments, in a manner that was highly reminiscent from previous works assessing goal-directed behaviours (Alcaraz et al., 2018; Parnaudeau et al., 2015). This pattern of results points toward a major role for the MD when only current mental representations can guide behaviour (Wolff et al., 2021; Wolff and Vann, 2019).

As a possible caveat, we cannot totally exclude a global performance issue in rats harbouring MD lesions as they appeared to constantly exhibit reduced responding during both the initial acquisition (Experiment 1) and the critical tests conducted under extinction conditions. This pattern of results indeed somewhat mirrors previous observations that rats with MD dysfunctions exhibit lower levels of instrumental performance during initial acquisition (Alcaraz et al., 2018; Corbit et al., 2003). Nonetheless, rats with MD lesions also exhibit normal levels of instrumental performance when environmental cues govern choice, suggesting that the deficit displayed by MD rats is at the associative - rather than performance – level (Ostlund and Balleine, 2008). Motivation *per se* is not affected by MD lesions as previously shown during either rewarded tests (Corbit et al., 2003), post-test reward consumption (Alcaraz et al., 2016; 2018) or progressive ratio assays (Alcaraz et al., 2016). Altogether, it thus seems that a performance deficit alone cannot satisfactory explain the current set of results. Additionally, as our lesions targeted the whole MD region, well beyond the central segment that more specifically connects to the OFC (Alcaraz et al., 2016; Mitchell and Chakraborty, 2013), the value of any functional inferences at circuit-level remains limited as dysconnectivity with multiple prefrontal areas may possibly account for the deficit exhibited by rats with MD lesions. It is however worth reminding that while the OFC is clearly associated with flexible behaviors based on stimulus-outcome association, the medial prefrontal cortex is not (Corbit and Balleine, 2003; Coutureau et al., 2012).

The present study thus complements earlier findings showing that both the OFC and the submedius thalamus are necessary to support successful performance during Pavlovian degradation (Alcaraz et al., 2015). While the role of the OFC has been well established by multiple labs (Balleine et al., 2011; Delamater, 2007; Ostlund and Balleine, 2007), the existence of convergent inputs from distinct thalamic nuclei may be one key functional aspect of this integrative function. We previously uncovered the role of the little known thalamic submedius nucleus connecting to the OFC (Alcaraz et al., 2015; Fresno et al., 2019) but surprisingly, the role of the other major thalamic input from the MD region has not been thoroughly examined in the same settings, with only mixed or indirect evidence thus far (e.g. Ostlund & Balleine, 2008; Pickens, 2008). The present study thus provides new insights on the role of the MD in a task that has previously been demonstrated to recruit both OFC and Sub functions. The current results indeed suggest that the MD may support dissociable functions with a possible initial role during acquisition of new Pavlovian associations (Experiment 1) and a consistent role in guiding behaviour when sensory feedback is no longer available to support adaptive responding after manipulating stimulus-outcome contingency (Experiments 1 & 2). While the former is highly reminiscent from multiple lesions studies showing that MD lesions typically slow down initial acquisition without necessarily preventing new learning in a wide range of behavioural tasks, even beyond Pavlovian learning (Chakraborty et al., 2019; Ouhaz et al., 2022; Wolff et al., 2015a, 2015b), the latter appears to be in line with other works showing that adaptive instrumental responding is also impaired when only current mental representations can guide performance (Alcaraz et al., 2018; Wicker et al., 2018).

As MD afferents are not the only source of thalamic inputs to the OFC, this questions the specific roles played by MD and Sub inputs and the necessity to have convergent sites within the OFC that can integrate both streams. One key difference between the Sub and the MD is that while they both project to the same OFC loci, only the latter also projects to other prefrontal areas such as the medial prefrontal cortex (Alcaraz et al., 2016). This suggests a more general role for the MD region, which may be more difficult to pinpoint than Sub functions. At first glance the role of the Sub indeed appears to be possibly more straightforward as Sub lesions totally spared Pavlovian acquisition but impaired degradation training in addition to the subsequent test under extinction conditions, suggesting a role in updating stimulus-outcome associations (Alcaraz et al., 2015). Interestingly, disconnecting the

Sub from the OFC also produced a specific impairment when rats were required to update current goal value after reversal of action and outcome identity, which was not observed when disconnecting the OFC from the MD (Fresno et al., 2019). Altogether, these data thus consistently support the idea that the Sub may play a specific role in updating associations that currently govern behavioral output. By contrast, the role of the MD does not seem to be the updating of current knowledge but rather to sustain or influence current mental representation to support performance when flexible responding is needed (Wolff et al., 2021; Wolff and Vann, 2019). The respective contributions of these thalamic nuclei is thus expected to be complementary rather than overlapping or competing. These functional considerations have possible relevance to further highlight the highly integrative role played by the OFC (Banerjee et al., 2020; Groman et al., 2019; Wang and Kahnt, 2021). Gaining a more systematic understanding of returning corticothalamic projections may be key to further advance our understanding of the functional principles at play within the thalamocortical architecture (Alcaraz et al., 2018; Choi et al., 2022; Harris et al., 2019).

In conclusion, the present paper provides new evidence that the MD critically supports performance when only recently updated stimulus-outcome associations can guide choice. Two independent experiments indeed confirmed a selective impairment when the sensory feedback provided by food outcome was not available to the animals, irrespective of whether or not initial acquisition was affected. Both experiments also suggested that updating stimulus-outcome associations was not prevented by thalamic damage, unlike the outcome of Sub lesions, consistent with a general role of the MD in guiding choices when animals must infer from the current associative framework. This general role of the MD supports the idea that this thalamic region may act as an important hub for cognitive functions which may be particularly affected in several mental conditions such as Schizophrenia (Anticevic et al., 2014; Parnaudeau et al., 2018) or drug addiction (Balleine et al., 2015; Huang et al., 2018).

## Acknowledgements

We thank Alain R Marchand for helpful discussion and analytic tools. Sarah Morceau was awarded a PhD grant EUR 17-EURE-0028 from the Bordeaux Neurocampus Graduate program.

